# Listening to Your Own Brain Waves Sound Enhances Your Sleep Quality and Quantity

**DOI:** 10.1101/2025.03.31.646309

**Authors:** A. Aloulou, M. Chauvineau, A. Destexhe, D. Léger

## Abstract

This pilot study examined the effects of relaxing personalized sound sequences (PSS), derived from individual slow-wave brain activity on sleep in adults with subjective insomnia complaints. Thirteen participants underwent one-night polysomnography to record delta wave activity (0.5–4 Hz), which was then transformed into individualized sound sequences. A randomized, single-blind, crossover protocol was then conducted at home, including two conditions of 3 to 5 consecutive nights: listening to the PSS and a non-personalized placebo sound sequence (PLA) for 30 minutes at bedtime. Objective sleep was assessed using a dry-electroencephalographic (EEG) headband and subjective sleep with a digital sleep diary. Compared to PLA, the PSS condition significantly increased total sleep time (Δ = +18.9 min, p = 0.05) and REM sleep proportion (Δ = +2.3%, p < 0.05), reduced REM latency (Δ = -16.6 min, p < 0.05) and improved overall sleep quality score (Δ = +1.4 A.U., p < 0.05). Participants with the shorter sleep duration (< 390 min, n = 5) and longer sleep onset latencies (> 20 min, n = 4) in PLA condition experienced greater improvements with PSS. These preliminary results suggest that listening to one’s own slow brain waves converted into sound may improve both sleep quality and quantity in individuals with moderate insomnia, with potentially enhanced benefits for those with more severe sleep difficulties.

**Highlights:**

- Personalized soundtracks were created by transforming each participant’s N3 slow-wave activity into audio sequences to listen to before bedtime.
- Participants reported significantly better subjective sleep quality when listening to their own brainwave-based soundtracks compared to placebo.
- Objective measures increased total sleep time and REM sleep during nights with personalized sound exposure.

## 1. Introduction

Sleep is a biological necessity essential for health, daily functioning and social development. Nevertheless, this process remains fragile, especially among individuals with chronic insomnia, who often struggle to achieve restorative sleep, leading to negative daytime consequences affecting well-being, health and social interactions [1]. Cognitive-behavioral therapy for insomnia (CBT-I) is currently the recommended first-line treatment for chronic insomnia, but it requires a rigorous and time-intensive protocol, and is not readily accessible in all settings [2]. On the other hand, pharmacological treatments, while commonly prescribed, may induce potential side effects and often fail to address the root causes of sleep disturbances [3].

In recent years, several non-pharmacological interventions using personalized approaches have emerged as promising alternatives to conventional insomnia treatments [4–6]. Among these, listening to relaxing music at bedtime has been suggested as an easy-to-implement and low-cost intervention to improve sleep quality and reduce insomnia symptoms [7–9]. Building on this concept, we investigated whether listening to sound sequences derived from each individual’s own slow-wave brain activity could provide measurable sleep benefits in people with moderate insomnia. If effective, this approach could contribute to the development of novel, personalized non-pharmacological interventions targeting sleep quality improvement in individuals with insomnia.

## 2. Materials and methods

### 2.1 Participants

Thirteen participants (7 females and 6 males; mean ± SD; age: 43.2 ± 11.0 years) with subjective complaints of insomnia participated in the study. Inclusion criteria, assessed through a medical examination, required age between 18 and 60 years, a poor perceived sleep quality (Pittsburgh Sleep Quality Index [10] score: 6-15), and absence of other polysomnography-confirmed sleep disorders (i.e., sleep apnea, periodic limb movement syndrome, hypersomnia, circadian sleep rhythm disorders, narcolepsy). Night and shift workers were excluded. Personal data were protected under French CNIL regulations, and participants received compensation upon study completion. The study was conducted according to the Declaration of Helsinki (1964: revised in 2001), and the protocol was approved by the local ethics committee (CPP IDF II ; CPP 2018-05-06). All participants provided written informed consent.

### 2.2 Experimental Design

The protocol included one in-lab inclusion night and two home-based experimental conditions, each lasting three to five nights in a randomized, single-blinded crossover design: one with a personalized sound sequence (PSS) and one with a placebo sequence (PLA). The inclusion night included a polysomnography recording to confirm the inclusion criteria and collect electroencephalogram (EEG) signals for customizing sound sequences. Before the protocol, each participant selected one of three versions of their PSS. The PLA condition used a non-personalized version of the selected sequence. Both versions were relaxing, and participants were blind to the conditions, unaware of their differences. Each night, the sound sequences were listened to using a wireless music headband worn at bedtime, with the sound automatically stopping after 30 min. A portable EEG headband was worn to monitor sleep, and an electronic sleep diary was completed upon waking. Participants were instructed to maintain usual diet, sleep habits and environment.

### 2.3 EEG Portable Headband

Objective sleep was recorded using a reduced-montage dry-EEG headband device (Dreem 3, Dreem, France). Participants were instructed to place the headband just before going to bed and remove it following get-up time. This device is a user-friendly alternative to polysomnography for automatically detecting waking states and sleep stages (light sleep = N1 + N2, SWS, REM) in 30-s epochs from brain activity via five EEG dry electrodes (F7, F8, Fp1, O1, O2). Automatic sleep-staging classification has been previously validated against polysomnography for sleep staging, achieving an overall accuracy (83.5 ± 6.4%) comparable to that of sleep experts [11]. Data were transmitted daily, enabling real-time issue resolution. Recordings containing fewer than 80% of “off-head” 30-s epochs were excluded.

### 2.4 Subjective Sleep Ratings

Each morning, participants completed the Spiegel Sleep Inventory (SSI) [12], a six-item self-administered questionnaire evaluating sleep initiation, quality, duration, night-time awakenings, dreaming and morning refreshment. Each item is rated from 1 to 5, with total scores ranging from 6 to 30.

### 2.5 Sound Conversion Method

A previous method was proposed to convert slow-wave brain activity into sound based on a parametric approach [13]. In this study, we employed a different conversion method, which consists of fitting the EEG signal corresponding to delta brain waves (0.5–4 Hz), recorded during the SWS periods in the inclusion polysomnography, with a sum of Gaussian functions. These functions were then used to modulate the amplitude of predefined digital sound samples. This approach automatically transforms the temporal structure of the EEG signal into a continuous and smooth sound sequence that mirrors the original brainwave patterns. Specifying the range of widths of the Gaussians allows us to automatically select the type of EEG waves encoded. Widths in the range of 100 to 1000 ms were used to focus on delta-frequency activity. Each generated Gaussian sequence was manually checked to ensure accurate encoding of the EEG signal. The resulting sound sequence was looped to produce a 30-minute track, with a linear fade-out applied so that the volume gradually decreased to silence by the end of the sequence. In the PLA condition, the same digital samples were used, but the Gaussian parameters—amplitude, width, and temporal position—were randomly shuffled to eliminate any direct correspondence with the participant’s EEG.

### 2.6 Statistical Analysis

Sleep data were compared across conditions using linear mixed-effects models (“lme4” R package, R Core Team, version 4.4.2), with participant as a random effect and condition as a fixed effect. Residual normality was tested using the Shapiro–Wilk test (p < 0.05). When violated, a 1,000-iteration bootstrap was applied to estimate 95% confidence intervals (CIs). Statistical significance was set at p < 0.05 (Wald tests, Kenward–Roger degrees of freedom). Results are reported as mean ± standard deviation (SD).

## 3. Results

A total of 93 out of 104 possible recordings (i.e., 89.4%) from the EEG headband were included in the analysis (mean ± SD nights per participant, PSS = 3.6 ± 1.0, PLA = 3.5 ± 0.8, p = 0.94). A slight delay in lights-on (Δ = +24.6 min, p < 0.05) was noted, along with increases in TST (Δ = +18.9 min, p = 0.05) and the proportion of REM (Δ = +2.3%, p < 0.05), and a reduction in REM latency (Δ = -16.6 min, p < 0.05) in PSS compared to PLA (Table 1). Perceived sleep duration (Δ = +0.2 A.U., p < 0.05), night-time awakenings (Δ = +0.3 A.U., p < 0.05), morning refreshment (Δ = +0.3 A.U., p < 0.05) and overall SSI score (Δ = +1.4 A.U., p < 0.05) were significantly improved in PSS compared to PLA. None of the other sleep data were different between conditions (p > 0.08, Table 1). Visual inspection of Figure 1 suggests that individuals with shorter TST (< 390 min, n = 5) and longer SOL (> 20 min, n = 4) in PLA exhibited a greater improvement in TST and reductions in SOL under the PSS condition, compared to those with longer TST (> 390 min, n = 8) and shorter SOL (< 20 min, n = 9).

**Table 1.** Objective and subjective sleep variables for the personalized sound (PSS) and the placebo (PLA) condition (mean ± SD).

|  | PLA | PSS |
| --- | --- | --- |
| **EEG headband** |  |  |
| Lights-off (h:min) | 23:54 ± 00:47 | 00:06 ± 01:01 |
| Lights-on (h:min) | 07:36 ± 00:59 | 08:00 ± 01:15* |
| Time in bed (min) | 462.4 ± 52.9 | 474.0 ± 65.0 |
| Total sleep time (min) | 395.6 ± 64.2 | 414.5 ± 67.6* |
| Sleep onset latency (min) | 22.0 ± 21.5 | 19.0 ± 22.1 |
| Wake after sleep onset (min) | 38.5 ± 34.8 | 34.5 ± 34.3 |
| Awakenings (n) | 22.2 ± 8.6 | 22.9 ± 8.4 |
| Sleep efficiency (%) | 85.6 ± 10.5 | 87.5 ± 8.4 |
| REM latency (min) | 88.6 ± 40.2 | 72.0 ± 30.6* |
| Light sleep (%) | 55.2 ± 12.6 | 53.9 ± 13.0 |
| SWS (%) | 18.5 ± 9.0 | 16.4 ± 7.8 |
| REM sleep (%) | 26.7 ± 6.9 | 29.0 ± 8.4* |
| Cortical arousals (n/h) | 12.4 ± 5.8 | 11.3 ± 5.6 |
| **Subjective sleep (SSI)** |  |  |
| Sleep initiation (A.U.) | 3.0 ± 1.0 | 3.3 ± 1.1 |
| Sleep quality (A.U.) | 2.8 ± 1.0 | 3.1 ± 0.9 |
| Sleep duration (A.U.) | 2.8 ± 0.7 | 3.0 ± 0.7* |
| Awakenings (A.U.) | 3.4 ± 0.8 | 3.8 ± 0.7* |
| Dreams (A.U.) | 4.2 ± 0.9 | 4.1 ± 0.8 |
| Feeling refreshed in the morning (A.U.) | 2.7 ± 0.8 | 3.0 ± 0.9* |
| Overall SSI score (A.U.) | 18.9 ± 3.5 | 20.3 ± 3.1* |

*Abbreviations:* REM, rapid-eye movement (REM); SWS, slow-wave sleep; SSI, Spiegel Sleep Inventory; A.U., arbitrary unit. *Note:* \*significantly different at $p < 0.05$ .

**Figure 1.**
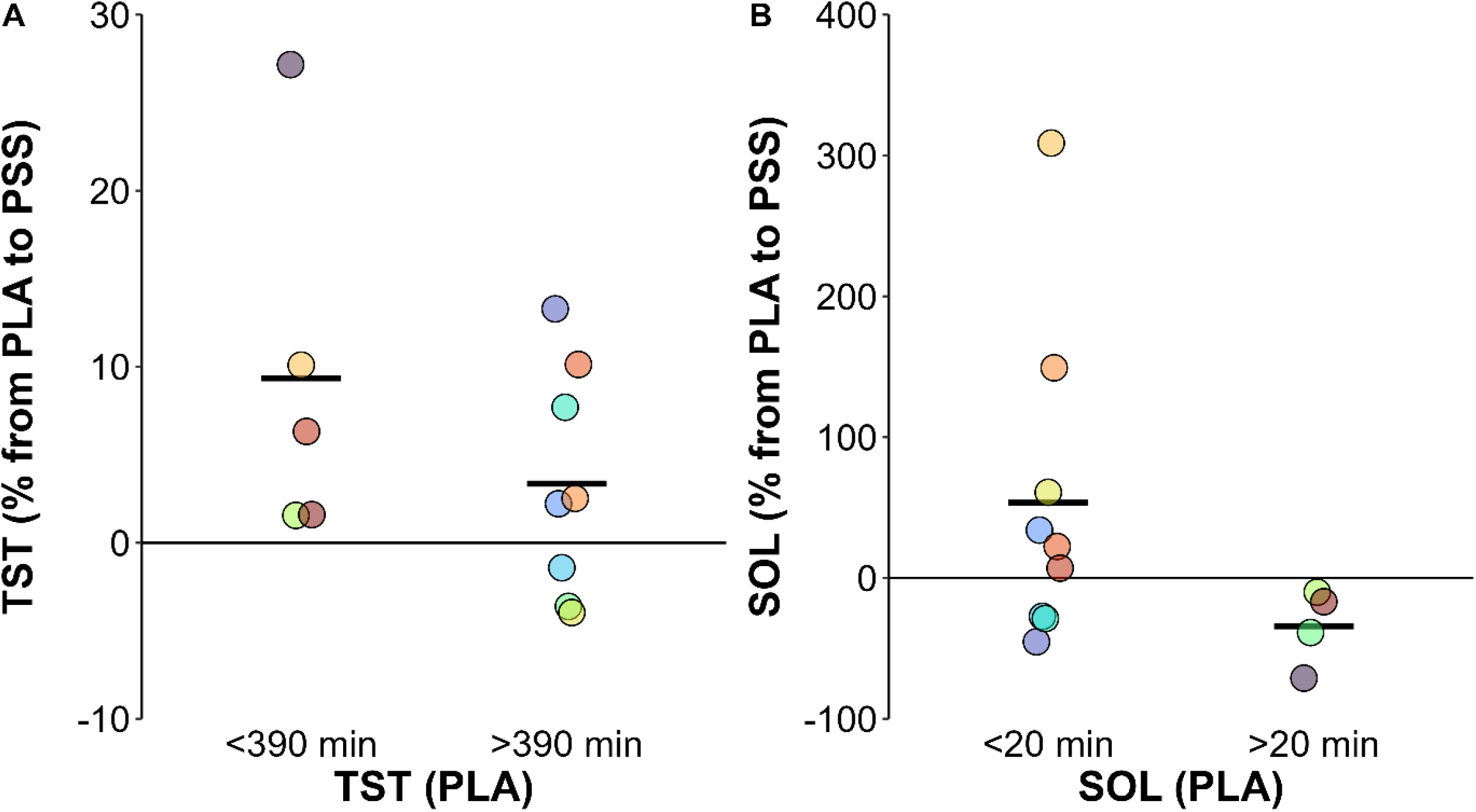
Magnitude of change in (A) total sleep time (TST) and (B) sleep onset latency (SOL) in the personalized sound (PSS) condition relative to the placebo (PLA) condition. *Notes*: Horizontal lines are means and each dot is an individual.

## 4. Discussion

The study provides novel evidence supporting the benefits of listening to one’s own slow-wave brain activity for improving sleep in individuals with insomnia. Participants slept 18.9 min longer in the PSS condition compared to PLA, and reported improvements in perceived sleep duration, sleep continuity and morning refreshment, resulting in significantly higher global sleep quality scores [12]. Notably, all participants who displayed the shortest TST (< 390 min) and longest SOL (> 20 min) under PLA showed greater improvements under PSS, suggesting that PSS may be particularly effective in shorter sleepers or those with sleep initiation difficulties.

Previous research has highlighted the effects of specific auditory stimuli, such as pink noise, on enhancing slow-wave sleep (SWS) [14]. Given that the sleeping brain can differentially respond to white noise across sleep stages, it is plausible that it may also respond selectively to sounds sharing temporal characteristics with delta waves [15]. Although PSS did not significantly affect SWS duration in this study, it was associated with a meaningful modulation of rapid eye movement (REM) sleep. Participants experienced an 8.6% relative increase in REM proportion and a 16.6-minute (18.7%) reduction in REM latency under PSS compared to PLA. These findings suggest that listening to brain-derived auditory patterns may facilitate earlier and more abundant REM sleep, a stage essential for cognitive functioning and emotional regulation [16].

Some limitations need to be considered when interpreting the present findings. While all participants reported subjective insomnia symptoms, most presented objective sleep values within normative ranges [17]. Only three participants had an average SOL > 30 min, five had WASO > 40 min, one had a sleep efficiency < 75%. Nine participants experienced insufficient sleep (< 7 h) [18]. Although this was a field-based pilot study, measures were taken to ensure consistency between conditions: participants were instructed to maintain stable routines throughout the two-week protocol, including regular timing of sleep, meals, physical activity, and avoidance of naps. Weekends were excluded to minimize potential circadian disruptions. Nevertheless, some uncontrolled environmental and behavioral factors—such as caffeine intake, evening screen exposure, ambient noise, or the presence of a bed partner—may have influenced sleep outcomes.. Despite these limitations, the observed improvements in both objective and subjective sleep parameters support the potential of personalized sound interventions as a non-pharmacological strategy for improving sleep. These findings underscore the importance of considering individual neurobiological characteristics in sleep interventions [19]. Further research is warranted to explore the effect of listening to one’s own slow-wave brain activity in individuals with more severe chronic insomnia, to better understand the underlying mechanisms, and to assess long-term impacts on sleep architecture and well-being.

## Acknowledgments

The authors are grateful to all the participants for their effort and involvement in this study.

## Author contributions statement

AA, AD and DL conceived and designed the research. AA conducted the experiments. AA and MC analysed and interpreted the data. AA, MC and AD wrote the first draft of the manuscript. All authors contributed to writing the final paper and approved the submitted version.

